# Phytohormones regulate asexual *Toxoplasma gondii* replication

**DOI:** 10.1101/2023.09.05.556389

**Authors:** Tina Wagner, Berit Bangoura, Stefanie Wiedmer, Arwid Daugschies, Ildiko Rita Dunay

## Abstract

The protozoan *Toxoplasma (T*.*) gondii* is a zoonotic disease agent causing systemic infection in warm-blooded intermediate hosts including humans. During acute infections, the parasite infects host cells and multiplies intracellularly in the asexual tachyzoite stage. In this stage of the life cycle, invasion, multiplication, and egress are most critical events in the parasite’s replication. *T. gondii* features diverse cell organelles to support these processes, including the so-called apicoplast, an endosymbiont-derived vestigial plastid originating from an alga ancestor. Previous studies have highlighted that phytohormones can modify the calcium-mediated secretion, e.g., of adhesins involved in parasite movement and cell invasion processes. The present study aimed to elucidate the influence of different plant hormones on the replication of asexual tachyzoites in a human foreskin fibroblast (HFF) host cell culture. *T. gondii* replication was measured by the determination of *T. gondii* DNA copies via qPCR. Three selected phytohormones, namely abscisic acid (ABA), gibberellic acid (GIBB), and kinetin (KIN)as representatives of different plant hormone groups were tested. Moreover, the influence of typical cell culture media components on the phytohormone effects was assessed. Our results indicate that ABA is able to induce a significant increase of *T. gondii* DNA copies in typical supplemented cell culture medium when applied in concentrations of 20 ng/μl or 2 ng/μl, respectively. In contrast, depending on the culture medium composition, GIBB may potentially serve as *T. gondii* growth inhibitor and may be further investigated as a potential treatment for toxoplasmosis.

## Introduction

The globally occurring parasite *Toxoplasma (T*.*) gondii* has a facultative two-host cycle (Tenter et al. 2000). It is assumed that all homoio- and also some poikilothermic animals including humans and livestock may serve as intermediate hosts (Dubey et al. 1998, Dlugonska et al. 2017), while only felines including domestic cats serve as definitive hosts where sexual recombination leads to fecal oocyst excretion (Dubey et al. 1970, Di Genova et al. 2019) Intermediate hosts are commonly infected orally by *T. gondii* oocysts via contamination of vegetables or other food with feline feces, or by tissue cysts contained in undercooked or raw meat (Jones and Dubey 2012). Another infection route for intermediate hosts includes transplacental infection of the offspring (Cowen and Wolf 1950, Dubey et al. 1995). Infective parasite stages include if ingested, the oocyst stage containing eight infective sporozoites each, if ingested, the tissue cysts containing up to hundreds of individual infective bradyzoites, and tachyzoites – the active asexual replication stage found in the intermediate host – mostly in the case of transplacental infections or e.g., blood transfusions with contaminated donor blood (Dubey et al. 1970, Dubey et al. 1998, Alvarado-Esquivel et al. 2018).

In intermediate hosts, after oral or transplacental infection with either of the named stages, the parasite undergoes asexual intracellular parasite replication supported by almost all nucleated cell types (Dubey et al. 1998). During acute infection, rapidly dividing stages (tachyzoites) multiply by endodyogeny. Under the influence of the host’s immune response, the tachyzoites later transform into slow-replicating permanent stages (bradyzoites in tissue cysts) (Watts et al. 2015). After crossing the blood-brain barrier, bradyzoites occur primarily in cells of the central nervous system (CNS) or in muscle cells though many other tissue types may be affected (Dubey et al. 1970, Dubey et al. 1998, Gagne 2001, Figueiredo et al. 2022). CNS persistence of bradyzoite-containing cysts results in specific immune responses and consecutive neuronal alterations (French et al. 2022, Steffen et al. 2022, Matta SK et al. 2018, Lang D et al. 2018, Parlog et al. 2015) These latent stages linger until an external stimulus like the host’s immunosuppression, or tissue cyst consumption and digestion by another host reactivates them. While little is known about the exact mechanisms of bradyzoite-tachyzoite interconversion (Tu et al. 2018), reconversion to the tachyzoite stage occurs with renewed invasion of host cells and intracellular proliferation (Dubey et al. 1998).

In *T. gondii* tachyzoites, movement and cell invasion are controlled by calcium-mediated secretion e.g., of adhesins to interact with host cell receptors during invasion (Chini et al. 2005). This process depends on the second messenger cADPR (cyclic adenosine diphosphate ribose), which activates various signal transduction and signal pathways (Moreno and Docampo 2003, Bothwell et al. 2005, Chini et al. 2005). It has been described that, cADPR controls microneme secretion and host cell invasion (Chini et al. 2005) as well as egress from host cells in *T. gondii* (Guse 2008). In turn, cADPR levels in *T. gondii* stages have been shown to be increased by abscisic acid (ABA), a plant hormone synthesized in endosymbiont organelle, the apicoplast (Guse 2008, Nagamune et al. 2008). The original function of ABA in plants is related to seed development, germination, vegetative growth, and stress tolerance (Sah et al. 2016), mediated by an increase in the concentration of cADPR, thereby stimulating the release of intracellular calcium (Wu et al. 1997, Puce et al. 2004, Zocchi et al. 2001). Besides ABA, there are several conserved plant hormones exhibiting partially antagonistic effects, regulating plant growth and development. Plant hormones include growth-promoting substances, such as auxins, cytokines, and gibberellins, or growth-limiting or otherwise regulating compounds, e.g., abscisic acid and ethylenes (Decker et al. 2006) In general, plant hormones are signal molecules with different chemical structures that are produced in plants and can influence physiological processes by promoting, suppressing or modifying them. (Tan et al. 2012).

Some recent studies suggest a positive therapeutic effect of plant hormones in mammalian systems (Trypuc et al. 2016). ABA has been shown to constitute an autocrine regulator for stem cell function with the potential to modulate inflammatory and immune responses (Scarfi et al. 2008). Phytohormones from the auxin group may be important as antitumor agents in human cells (Hamayun et al. 2017). Gibberellin was shown to exhibit an enhancing effect on cell division and apoptosis in toads (Sakr and Shalaby 2012). Since the 1940s, several therapeutics have been developed for *T. gondii* infections. However, the current gold standard, a combination of pyrimethamine and sulfadiazine (Pyr-Sulf) targeting the acute stage of infection still has significant failure rates (Eyles and Coleman 1953, Dunay et al. 2018). Hence, despite significant advances in the treatment of human toxoplasmosis, there is a strong impetus for ongoing development of new therapeutics for both acute and latent disease stages. In this regard, the present study may further help to explore new therapeutic approaches.

Based on prior studies that showed a direct effect of external ABA supplementation on cADPR levels and the known plant-derived origin of the apicoplast in *T. gondii* (Nagamune et al. 2008, McFadden 2011), the present *in vitro* study aimed to elucidate the influence of different plant hormones on the individual asexual stage tachyzoites with respect to their reproduction.

## Material and Methods

### Study design

Before testing the effect of three selected phytohormones ABA, gibberellic acid (GIBB), and kinetin (KIN)) on *T. gondii* tachyzoites, two pre-assays were performed (please see Fig. 1 for study design).

**Figure 1:**
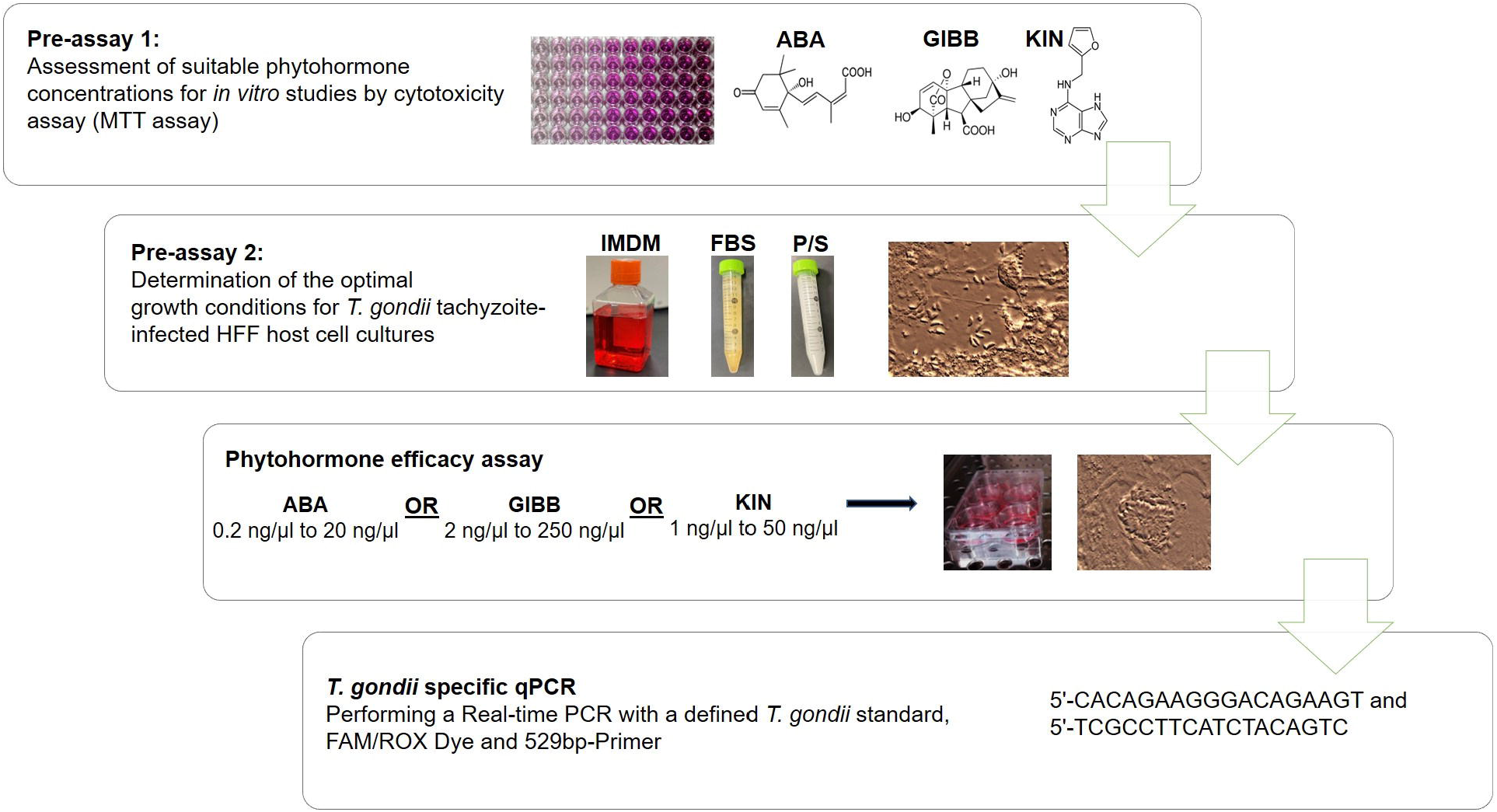
Experimental design; The schematic workflow shows the sequential steps of this study. MTT (3-(4,5-dimethylthiazol-2-yl)-2,5-diphenyltetrazolium bromide); ABA (abscisic acid); GIBB (gibberellic acid); KIN (kinetin). HFF (human foreskin fibroblast); IMDM (Iscove’s modified Dulbecco’s medium); P/S (penicillin/streptomycin); FBS (fetal bovine serum)

#### Pre-assay 1

A cytotoxicity assay (MTT assay) using the 3-(4,5-Dimethylthiazol-2-yl)-2,5-diphenyltetrazolium bromide dye (Mosmann 1983) was performed to evaluate the host cell culture tolerance towards different phytohormone concentrations.

#### Pre-assay 2

The optimal growth conditions for *T. gondii* tachyzoite-infected host cell cultures were determined (see Table 1). Therefore, human foreskin fibroblast (HFF, ATCC: SCRC-1041™) cell cultures infected with the *T. gondii* type II strain ME49 (ATCC® 50611, Wesel, Germany) were subjected to different growth media containing pure Iscove’s modified Dulbecco’s media (IMDM, Gibco, Invitrogen, Karlsruhe, Germany) or added penicillin/streptomycin (P/S, GE Healthcare, Germany) and/or fetal bovine serum (FBS), respectively to assess the replication rate of *T. gondii* under these varied conditions.

#### Actual phytohormone efficacy experiments

Based on the MTT assay results, *T. gondii* tachyzoite infected host cell cultures were treated with different ABA, GIBB, and KIN concentrations. At the same time, a possible influence of the above-mentioned medium growth additives (presence or absence of P/S and/or FBS) on the efficacy of the phytohormones was investigated (see Table 2). The effect of phytohormone-growth medium combinations on tachyzoite multiplication was assessed by quantifying the final *T. gondii* DNA copy number per well for each treatment group. The *T. gondii* yield was compared with an infected untreated control group (IUC).

### Parasites, parasite maintenance and infection

For the *in vitro* experiments, *T. gondii* type II strain ME49 (ATCC® 50611, Wesel, Germany) was used. In line with previous descriptions (Khan and Grigg 2017), the *T. gondii* strain was maintained in the continuously reproducing tachyzoite stage in HFF cultures passaged once to twice per week. Host cell cultures were grown at 37 °C and 5 % CO_2_ in Dulbecco’s Modified Eagle Medium (DMEM, Gibco, Invitrogen, Karlsruhe, Germany) supplemented with 10 % heat inactivated (at 56 °C for 45 min) FBS and 1 % P/S until approx. 80 % confluent for *T. gondii* infection. *T. gondii* infected HFF cultures were maintained in IMDM, that was supplemented with FBS and/or P/S depending on the experimental group during pre-assay 2 and the phytohormone efficacy experiments. Infected cultures were incubated as described above.

### Phytohormones

Three phytohormones (ABA, GIBB, and KIN) were tested. Stock solutions were prepared as follows: *ABA:* 10 mg ABA (Cayman chemicals cat. ID.: 10073) were dissolved in 1.25 ml 50 % dimethyl sulfoxide (DMSO, Sigma Aldrich Chemie GmbH, Taufkirchen, Germany) over 2 h at 40 °C while shaken. The resulting stock solution of 8 mg ABA /ml was sterile filtered, aliquoted, and stored at −20 °C until experimental use. *GIBB:* 5 mg GIBB powder (Carl Roth GmbH & Co. KG, Karlsruhe, Germany) was dissolved in 1 ml double distilled water (ddH_2_O) to obtain a stock solution of 5 mg GIBB /ml. After sterile filtration, the GIBB solution was aliquoted and stored at 4 °C in the dark. *KIN:* 21.5 mg KIN powder (Gold Biotechnology, St. Louis, MO, USA) was dissolved in 1 ml 1M NaOH, and ddH_2_O was added to the resulting solution to a total volume of 10.75 ml, i.e., the stock solution contained 2 mg KIN /ml. The stock solution was sterile filtered, aliquoted, and stored at 4 °C until use.

All stock solutions were diluted in IMDM, IMDM supplemented with 10 % FBS, or IMDM supplemented with 10 % FBS and 1 % P/S, respectively, for direct use on *T. gondii*-infected HFF cultures.

### Pre-assay 1: MTT assay

Briefly, 2 × 10^4^ HFF cells per well were seeded into 96-well plates and incubated for 24 hours (until a confluent monolayer was formed). Subsequently, 100 μl/well of the respective phytohormone solution was added. All phytohormones were diluted from their stock solutions (please see below) in DMEM Concentrations tested were 2,000 ng/μl, 400 ng/μl, 200 ng/μl, 20 ng/μl, 2 ng/μl, and 0.2 ng/μl for ABA; 1,250 ng/ml, 125 ng/μl, 12.5 ng/μl, 1.25 ng/μl, and 0.125 ng/ml for GIBB and 1,000 ng/ml, 500 ng/μl, 50 ng/μl, and 0.5 ng/ml for KIN, respectively. As a negative control, untreated cultures incubated in DMEM were used. After a 48-h incubation period, the MTT assay was performed as described previously (Thabet et al. 2015, Kumar et al. 2018) and cytotoxicity was assessed.

### Pre-assay 2: Assessment of optimal *T. gondii* replication conditions in the selected host cell culture system

Confluent HFF cultures in 6-well plates were infected with 5 × 10^5^ *T. gondii* tachyzoites per well. The infected cultures were incubated for 48 h after infection (hpi) with either pure IMDM or IMDM with supplementation of growth additives (see Table 1). At 48 hpi, cultures were detached and frozen for later DNA extraction and *T. gondii*-specific quantitative real-time PCR (qPCR) as outlined below. based on qPCR results, absolute *T. gondii* numbers were calculated for each experimental group as described below.

### Phytohormone efficacy experiments

In the main experiment, the effect of phytohormones on the proliferation of tachyzoites and a possible influence of the growth additives on this effect were investigated. For this purpose, either ABA, KIN, or GIBB were diluted in IMDM medium at different concentrations (see Table 2) and added to the HFF monolayers. Concentrations tested ranged from 0.2 to 20 ng/μl for ABA, from 2 to 250 ng/μl for GIBB, and from 1 to 50 ng/μl for KIN. Dilutions were prepared in different growth media variants, i.e., IMDM either with or without FBS and/or P/S, as outlined in Table 2. This was done to investigate any possible interactions between the phytohormones used and the growth additives FBS and P/S to evaluate if any findings were related to varying cell culture conditions. Directly after group-specific treatments, the cultures were infected with 5 × 10^5^ *T. gondii* tachyzoites. The medium containing the diluted phytohormones was removed after 2 h of incubation with the infected cultures and replaced by unmedicated cell culture medium. Therefore, the supernatant was removed and replaced for all groups identically with fresh, unmedicated growth medium IMDM + 10 % FBS + P/S, which was identified in pre-assay 2 as optimal growth medium to support *T. gondii* tachyzoite replication for the follow-up replication period of 48 h. The removed supernatant was centrifuged (1540 × g, 5 min) in a 1.5 ml microcentrifuge tube separately per each cell culture well to collect any free tachyzoites, and the pellet was resuspended in the unmedicated IMDM + 10 % FBS + P/S before being returned to each respective well. As infected untreated controls (IUC), groups of unmedicated wells in different growth media (either incubated for 2 h initially in IMDM, IMDM + 10 % FBS, IMDM + P/S, or IMDM + 10 % FBS + P/S, respectively, according to the media used in medicated groups for the 2 h phytohormone incubation) were carried along (see Table 2, all IUC groups). After an incubation period of 48 hpi, the cultures were detached using accutase cell dissociation reagent (Gibco, Invitrogen, Karlsruhe, Germany). Therefore, the growth medium was removed and collected and centrifuged per cell culture well as described above for the media change. The resulting pellet was combined with the pellet from accutase treatment of the adherent cell culture. Cultures were incubated with accutase solution for about 10 minutes (until cells were detached as confirmed by light microscopy). The accutase solution was removed by centrifugation (1540 × g for 5 min) and the resulting cell pellets were combined with the supernatant-derived pellets and re-suspended in 200 μl PBS (phosphate buffered saline) and stored at −20 °C until DNA extraction. The DNA extract for each individual cell culture well was analyzed quantitatively for *T. gondii* by qPCR as outlined below. This scheme was then followed by the investigation of a possible influence of growth additives (P/S and/or FBS) in the growth medium on the previously shown effect of the phytohormones (see Table 2 experimental groups A20-F through K1-PF).

#### DNA extraction and T. gondii quantitative real time PCR (qPCR)

DNA was extracted from each well contents with the QIAamp DNA Mini Kit (Qiagen, Hilden, Germany) following the instructions of the manufacturer.

Real-time PCR was performed using a defined *T. gondii* standard (extracted DNA from 1×10^6^, 5×10^5^, 1×10^5^, 5×10^5^, 1×10^4^, 1×10^3^, and 1×10^2^ ME49 tachyzoites, respectively) for product quantification

The 20 μl/reaction master mix contained 0.9 μM of each 529-bp gene fragment specific forward and reverse primer (5’-CACAGAAGGGACAGAAGT and 5’-TCGCCTTCATCTACAGTC) (Edvinsson et al. 2006), 12.5 μl Maxima Probe/ROX qPCR Master Mix (2X) (order no. K0231, Thermo Fisher Scientific, Germany), and nuclease free water to fill up the volume to 20 μl total volume. The total reaction volume was 25 μl after addition of 5 μl template DNA sample. The PCR was performed on a Stratagene MX3000P thermocycler (LaJolla, USA) using the following cycling program for all experiments: 1 cycle at 95 °C for 15 sec, followed by 45 cycles with 95 °C for 15 sec, 60 °C for 60 sec, and 72 °C for 15 sec. Resulting Ct values were calculated into DNA copy numbers based on the standard curve created from the tachyzoite standard concentrations given above.

#### Statistics

Data was analyzed descriptively using mean values and standard error of the mean (SEM), as well as median values and quartiles. All statistical data analyses were performed using the IBM SPSS Statistics for Windows software package (version 28.0, 2021, IBM Corp, Armonk, NY, USA). Data for pre-assay 2 experiments and phytohormone efficacy experiments were tested for normal distribution by Kolmogorov-Smirnov normality test. Since all data were non-normally distributed, group comparisons were performed using the nonparametric Mann-Whitney test. The IUC wells were used as comparative reference group. Statistical significance between study groups was defined at a level of p < 0.05.

## Results

### Pre-assay 1 (MTT assay)

In all phytohormone-treated cultures, the cell viability ranged from 81.4 to 96.5 % which was comparable to the untreated control cultures with 84.4 to 96.6 % viability. The cell viability ranged from 83.2 to 97.1 % for different solvents. Cell viability percentages of at least 80 % were considered suitable for further experimentation. Therefore, pre-assay 1 revealed that all tested concentrations of all three phytohormones and their respective solvents were tolerated well by the host cell cultures, i.e., no phytohormone or solvent-related cytotoxicity was observed (data not shown).

### Pre-assay 2 (Assessment of optimal *T. gondii* replication conditions in the selected host cell culture system)

In comparison to pure IMDM, a significant increase of *T. gondii* DNA copies were detected in all supplemented IMDM media (see Fig. 2). Parasite replication over the 48 hpi observance period was particularly high in the medium variant IMDM+FBS (with a mean *T. gondii* DNA copy number increase of 83.6 %) and in IMDM+10 % FBS + P/S (with *T. gondii* DNA copy numbers being increased by a mean value of 57.3 %). A significant replication-enhancing effect of FBS on *T. gondii* tachyzoites could be observed in the selected HFF culture model (p < 0.05). Based on these results, IMDM supplemented with 10 % FBS and P/S was considered most suitable for the 48 h incubation period in the phytohormone efficacy experiments to observe optimal post-treatment *T. gondii* tachyzoite replication under identical conditions for all groups.

**Figure 2:**
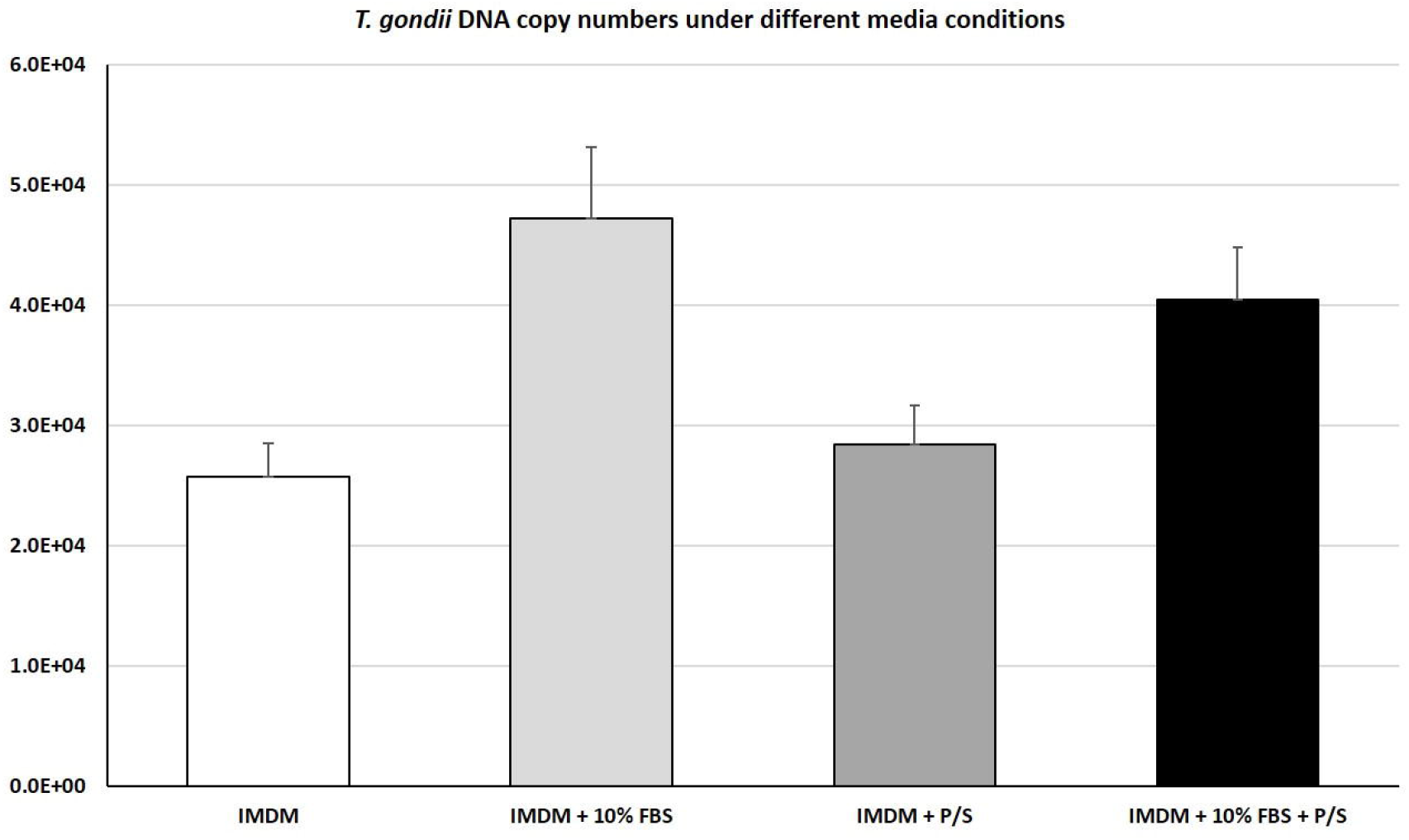
Pre-assay 2; Replication of *T. gondii* ME49 tachyzoites using different cell culture growth media compositions (mean values, error bars indicate standard error of the mean, SEM; n = 36) using different cell culture media compositions (IMDM, Iscove’s modified Dulbecco’s medium; P/S, penicillin/streptomycin; FBS, fetal bovine serum).

### Phytohormone efficacy experiments

The tested phytohormones exhibited varying effects on *T. gondii* tachyzoite replication (see Figs. 3 to 5), depending not only on the phytohormone used but also on the growth media applied (either pure IMDM or IMDM with FBS and/or P/S).

**Figure 3:**
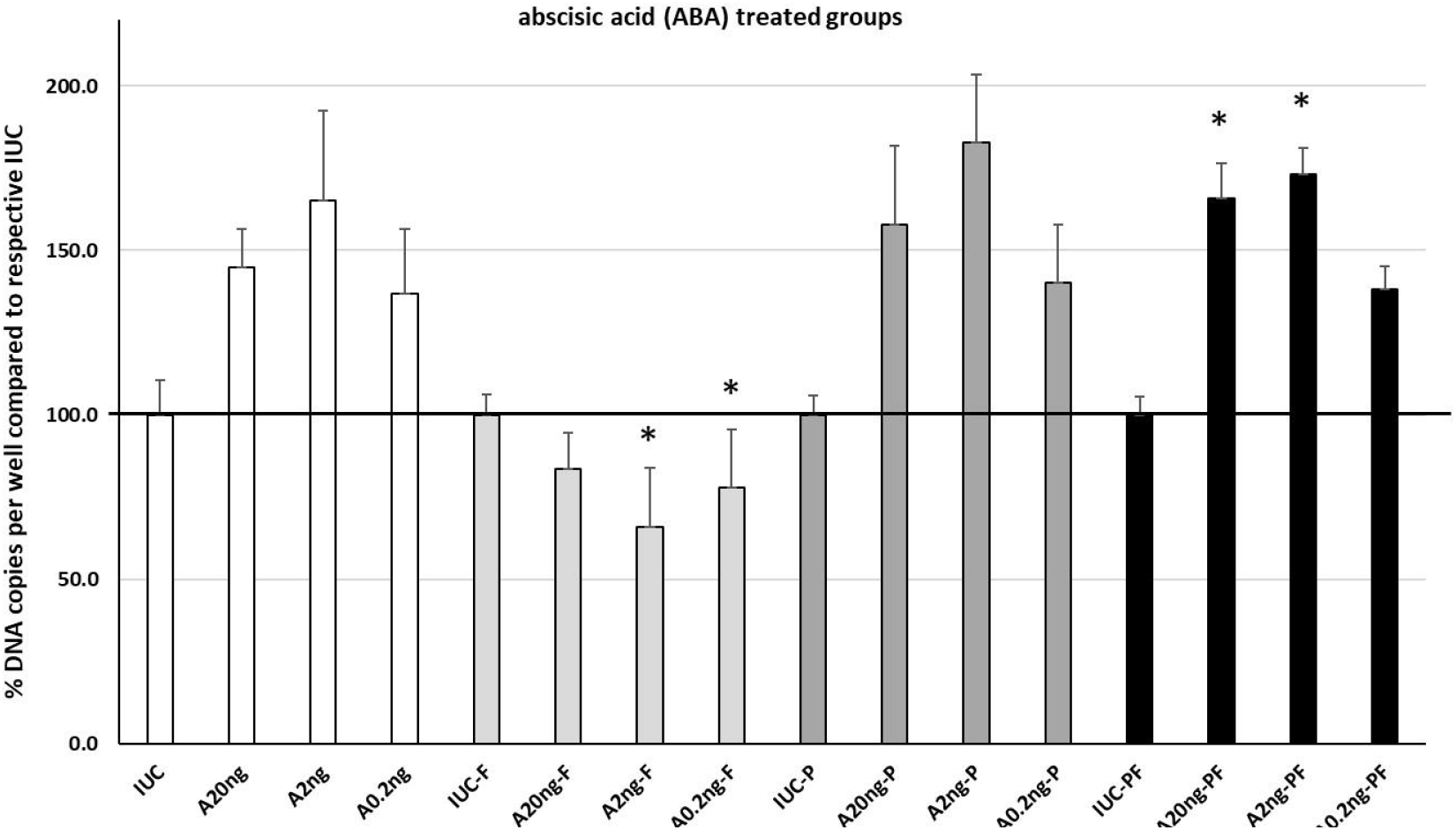
Effect of abscisic acid (ABA) on replication of *T. gondii* ME49 tachyzoites using IMDM with different cell culture conditions compositions (IMDM, Iscove’s modified Dulbecco’s medium; P, penicillin/streptomycin; F, fetal bovine serum) enriched with different ABA concentrations (mean values displayed as % of IUC (infected untreated control group) mean for the respective growth medium composition, error bars indicate standard error of the mean (SEM); n= 6 to 12). * indicates statistically significant difference (p < 0.05) compared with the IUC group of the respective media composition. For full explanations of the individual group abbreviations, please see table 2.

#### ABA

The phytohormone ABA (see Fig. 3) promoted the replication of *T. gondii* at all tested concentrations for all growth media except for IMDM + 10 % FBS used for groups A20ng-F, A2ng-F, and A0.2ng-F. Interestingly, the highest *T. gondii* DNA copy number increase compared to the respective IUC group could be observed consistently in the cell cultures treated with 2 ng ABA /μl, i.e. the medium ABA test concentration with mean values of 165.0 % of mean IUC (group A2ng), 182.9 % (group A2ng-F), and 173.0 % (group A2ng-PF), respectively. Regarding the *T. gondii* DNA copy number reduction observed for ABA in IMDM + 10 % FBS, it was also most pronounced in the 2 ng ABA /μl treated group A2ng-F with a mean reduction of 34.1 % compared to group IUC-F. A statistically significant effect could be observed for groups A2ng-F and A0.2ng-F, which showed inhibited parasite replication, and groups A20ng-PF and A2ng-PF, displaying enhanced parasite replication, respectively.

#### GIBB

In our experiments, GIBB suppressed the parasite replication consistently, with the slight exception of group G10ng-PF (see Fig. 4). The most pronounced inhibitory effects were seen in GIBB treated groups maintained in FBS-free media (IMDM or IMDM + P/S). In groups G250ng through G2ng, statistically significant (p < 0.05) *T. gondii* growth inhibition mean values of 93.6 to 98.5 % of the IUC mean value were detected. No statistically significant differences were seen between any of the groups G250ng, G50ng, G10ng, or G2ng. While not statistically significant, *T. gondii* copy numbers were markedly reduced in groups G250ng-P through G2ng-P as well, with inhibitory effects reaching a maximum of 86.6 % in high-dose GIBB treated group G250ng-P. Addition of FBS to the growth media seemed to reduce the inhibitory effect for IMDM + 10% FBS and IMDM + 10% FBS + P/S treated groups. In groups G250ng-F through G2ng-F, inhibition varied from 46.8 % (group G250ng-F) to 54.1 % (group G50ng-F); no clear GIBB dose dependency could be established. In IMDM + 10% FBS + P/S incubated groups, only group G250ng-PF showed a medium *T. gondii* DNA copy reduction level of 28.3 %, while all other groups showed *T. gondii* DNA amounts similar to the untreated IUC-PF group.

**Figure 4:**
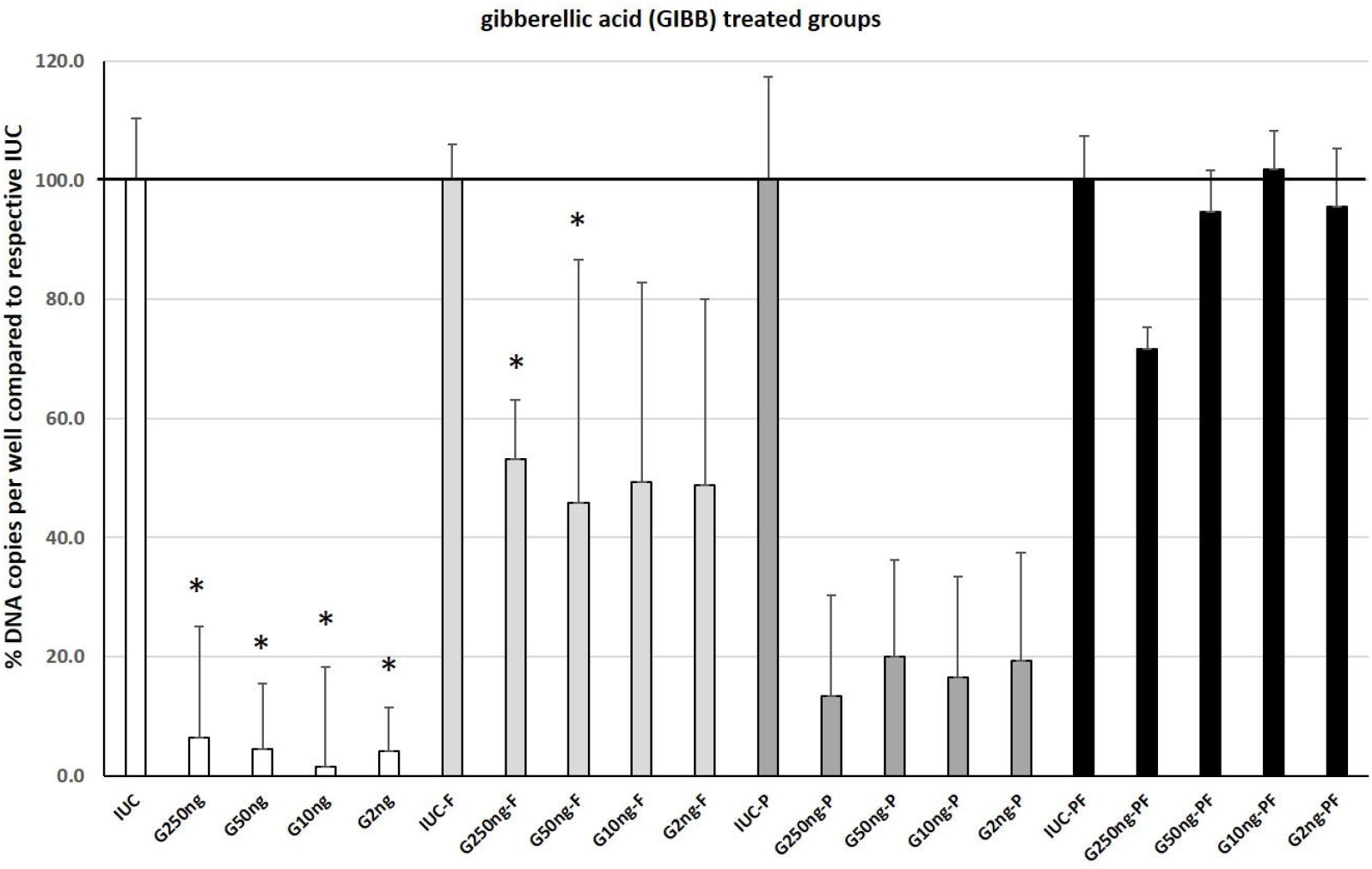
Effect of gibberellic acid (GIBB) on replication of *T. gondii* ME49 tachyzoites using IMDM with different cell culture conditions compositions (IMDM, Iscove’s modified Dulbecco’s medium; P, penicillin/streptomycin; F, fetal bovine serum) enriched with different GIBB concentrations (mean values displayed as % of IUC (infected untreated control group) mean for the respective growth medium composition, error bars indicate standard error of means (SEM); n= 6 to 12). * indicates statistically significant difference (p < 0.05) compared with the IUC group of the respective media composition. For full explanations of the individual group abbreviations, please see table 2.

#### KIN

The effect of KIN supplementation showed a strong variation, ranging from inhibition to promotion of *T. gondii* replication (see Fig. 5). In groups without P/S addition, the lower KIN test concentration groups displayed an increased parasite replication, with maximum values of 65.2 % (significant at p < 0.01) or 27.7 % increase observed in the low-dose treated groups K1ng-F or K1ng, respectively. For groups treated with IMDM or IMDM + 10 % FBS, there was an inversely dose-related KIN effect; the higher the KIN concentration applied, the lower was the obtained *T. gondii* parasite copy number. For groups K50ng-P, K25ng-P, and K1ng-P, there was a mildly positive effect on parasite replication. For groups treated with KIN in IMDM + 10 % FBS + P/S, a consistent decrease in parasite copy numbers was seen, with lowest parasite copy numbers in group K1ng-PF (26.2 % mean reduction compared to IUC-PF, significant at p < 0.05).

**Figure 5:**
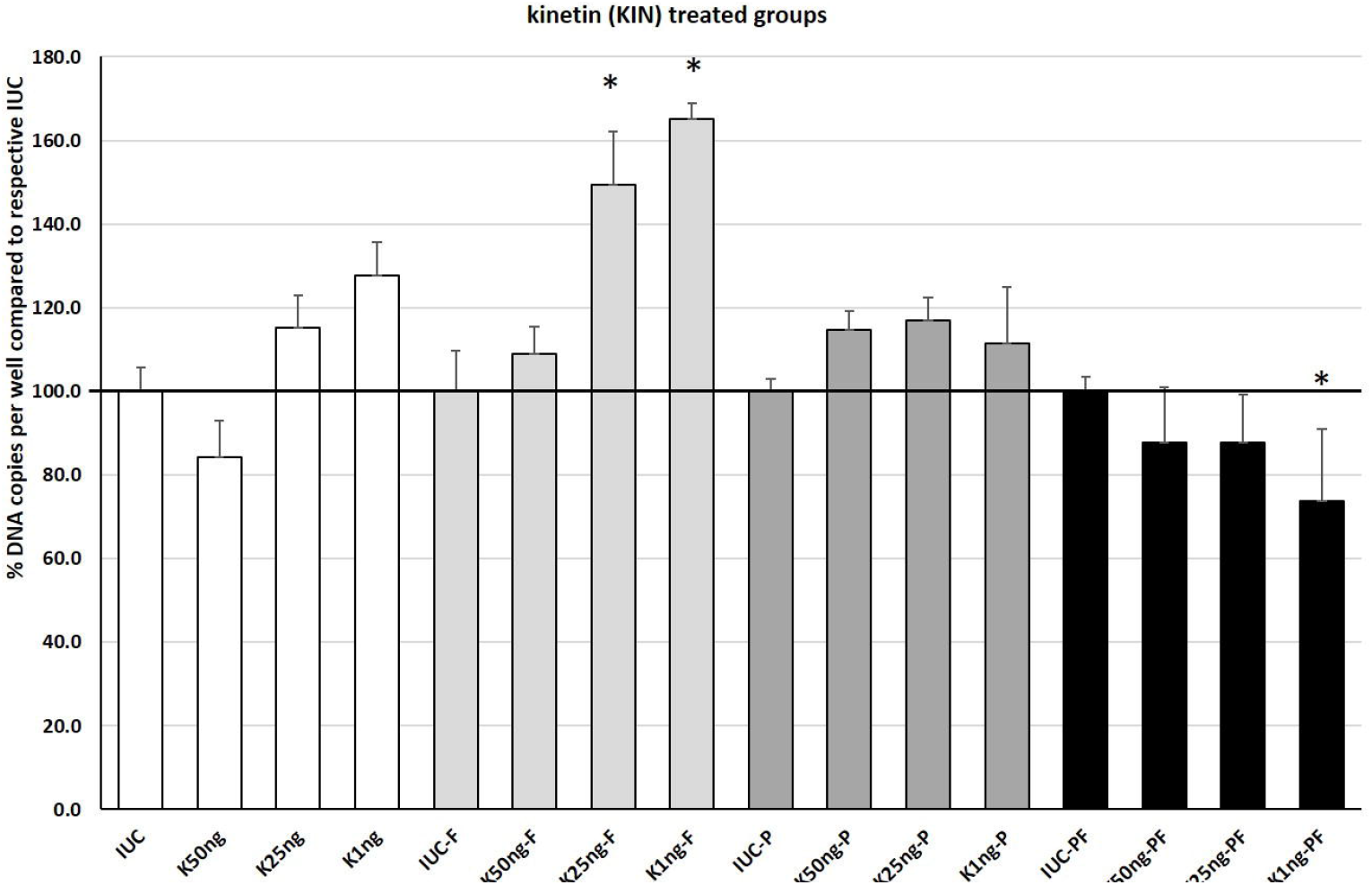
Effect of kinetin (KIN) on replication of *T. gondii* ME49 tachyzoites using IMDM with different cell culture conditions compositions (IMDM, Iscove’s modified Dulbecco’s medium; P, penicillin/streptomycin; F, fetal bovine serum) enriched with different KIN concentrations (mean values displayed as % of IUC (infected untreated control group) mean for the respective growth medium composition, error bars indicate standard error of the mean (SEM); n= 6 to 12). * indicates statistically significant difference (p < 0.05) compared with the IUC group of the respective media composition. For full explanations of the individual group abbreviations, please see table 2.

## Discussion

In the current study, we investigated the effect of three phytohormones (ABA, GIBB, and KIN) on the replication potential of *T. gondii* tachyzoites. *T. gondii* is an apicomplexan parasite known to harbor the algae-derived apicoplast as well as the plant-like vacuolar compartment (PLVAC) (Stasic et al. 2022). Therefore, it is assumed that plant-derived phytohormones may exhibit a significant effect on cell-signaling pathways related to organelles shared with plant organisms (Nagamune et al. 2008, Andrabi et al. 2018). Previous studies have indicated a direct effect of cytokinins, a class of phytohormones including KIN, on cell cycle progression as well as proliferation in *T. gondii* tachyzoites (Andrabi et al. 2018). Other studies implied that ABA is involved not only in Calcium-dependent intrinsic signaling pathways in *T. gondii*, but also in the parasite-induced host cell autophagy that supports parasite growth (Nagamune et al. 2008, Wang et al. 2009). Synthetic gibberellin inhibitors applied on *T. gondii in vitro* cultures have been shown to inhibit *T. gondii* growth (Toyama et al. 2012). The described studies indicate that phytohormones play a crucial role in regulating *T. gondii* tachyzoite replication, however, neither the direct comparison of different phytohormone classes in the same infection model nor the influence of other cell culture medium components such as widely used FBS and antibiotics have been addressed before. Our study aimed to establish the comparative effect of different previously described phytohormone groups (represented by ABA, GIBB, and KIN) in the same study under identical culture conditions as well as the effect of common media additives on the study outcome. We selected the three phytohormones based on their varying effects in plant growth and maturation as described above. In the literature, few information on utilized phytohormone concentrations in *T. gondii* cultures was available. For ABA, Nagamune et al. (2008) tested 0.0026 to 0.26 ng ABA /μl medium. Because of their highly significant finding for *T. gondii* cADPR production increase at their highest ABA test concentration, we selected concentrations encompassing and exceeding their test concentrations, i.e., 0.2 to 20 ng ABA /μl. Similarly, for cytokinins a dosage of 10 μM has been described (Andrabi et al. 2018), which translates into 2.2 ng/μl for KIN, falling within our tested range of 1 to 50 ng KIN /μl. For GIBB, we did not find information about previous concentrations used in the apicomplexan model, however, we referred to concentrations typically used in plant experiments and designed a test concentration range of 2 to 250 ng GIBB /μl, which is in line with plant experimentation (Didi et al. 2022, Santos et al. 2022).

In our experiments, we used the type II ME49 strain of *T. gondii*, which is a commonly used laboratory strain that has proven suitable for *in vitro* and *in vivo* studies, while it represents type II that is often encountered in human field infections (Hosseini et al. 2018).

In the selected model, we could see a response in *T. gondii* replication quantity to all phytohormones tested. Generally, ABA and KIN predominantly tended to promote *T. gondii* replication, while GIBB tended to exert a negative effect on parasite growth. However, these findings seemed to overlap with interactions with FBS and P/S that interfered with all three phytohormones. Phytohormones were left on the treated cell culture groups for 2 h in the varied growth media (IMDM with or without FBS and/or P/S). The post-treatment 48 h incubation was identical for all groups under the optimal *T. gondii* growth conditions as determined in pre-assay 2 (i.e., using IMDM + 10 % FBS + P/S). This was done to ensure that any group differences observed can be clearly attributed to any treatment effects during the phytohormone incubation rather than post-treatment growth conditions. Thus, any differences within a phytohormone concentration that are observed between different growth media utilized for phytohormone incubation are findings we attribute directly to an interaction between phytohormones and growth additives. For ABA, which promoted parasite growth under most conditions, intriguingly we observed a drop in *T. gondii* tachyzoite numbers only and consistently over all test concentrations when combined with IMDM + 10 % FBS, but not when P/S was added in. In *Arabidopsis* plants, an interaction between bovine serum albumin (BSA) added to culture media and root growth as mediated by phytohormones could be observed (Lonhienne et al. 2014). The authors observed a dose-dependent effect, with low amounts of BSA promoting root growth and high BSA concentrations inhibiting root growth. Based on the high BSA concentrations of roughly 50 mg/ml present in FBS (Francis et al. 2010), in our experiments, a high BSA concentration was present, that may potentially have interacted with ABA pathways and suppressed parasite growth potentially in line with the cited findings in *Arabidopsis* (Lonhienne et al. 2014). However, since our current study was not designed as mechanistic study, the underlying mechanism – for both the observed inhibitive effect of the ABA-FBS combination on *T. gondii* replication and its reversal when P/S was added – remains unclear. Any other of the tested media compositions lead to a promotion of parasite growth by ABA in any tested concentration. This is consistent with previous studies (Nagamune et al. 2008, Wang et al. 2009) that suggest a direct effect of ABA on cADPR-dependent parasite invasion and replication events. The negative effect of GIBB on parasite replication could be observed for any media conditions, however, it was most pronounced in the absence of FBS. It is known from plant studies that GIBB and ABA are partially antagonizing each other’s effects, mutually interacting to maintain plant homeostasis (Castro-Camba et al. 2022). Therefore, an antagonistic effect of both phytohormones on *T. gondii* was anticipated. Still, regarding apicomplexan parasites, our finding seems somewhat contradictive to literature descriptions of the detrimental effect of gibberellin inhibitors on *Plasmodium* (Toyama et al. 2012). Interestingly, Toyama et al. (2012) could not show a beneficial effect of external gibberellin addition to a *T. gondii* culture, although inhibition of endogenous gibberellins seemed to have an adverse effect on *T. gondii* replication. This indicates that adding plant GIBB may not lead to a resorption rate that leads to parasite growth promotion, or that plant gibberellins may be somewhat chemically varied from putative parasite gibberellins. Andrabi et al. (2018) could not detect endogenous GIBB production in apicomplexan parasites though inhibitor substances would impact the culture negatively, which again may indicate that there are plant-like but not identical phytohormones present in apicomplexan parasites. While currently, we cannot offer a substantiated explanation for the observed acute decline of parasite stage numbers in our GIBB treated groups, this is an interesting finding similar to GIBB effects on higher plants (Castro-Camba et al. 2022). For KIN, mixed effects on *T. gondii* growth were observed, seemingly dependent both on KIN concentration and on the selected cell culture media composition. In plants, KIN was discovered as a strong cell division promoter more than five decades ago already (Miller et al. 1956), and many more cytokinin functions have been added more recently (Werner et al. 2003). Andrabi et al. (2018) observed an increased copy number of apicoplasts per *T. gondii* tachyzoite in natural cytokinin-treated *T. gondii* cultures and could establish a regulatory effect of cytokinins on the parasite cell cycle progression. However, the authors stated partially contradictive effects for the different cytokines tested and suggested the possible existence of unidentified receptors for cytokinins in *T. gondii* (Andrabi et al. 2018). Regarding the effect on the cell cycle, ABA inhibitor studies indicate an involvement of ABA in stage conversion between tachyzoites and dormant bradyzoites (Nagamune et al. 2008). We may merely speculate that other phytohormones like GIBB or KIN may be involved in triggering stage conversion as well, potentially in opposite ways, which may potentially aid in explaining the low *T. gondii* replication specifically in GIBB treated groups but without more detailed knowledge on the phytohormone receptors and triggered biochemical reactions in *T. gondii* this currently remains unknown. Furthermore, we would like to suggest the future conduct of basic chemical studies into interactions between different phytohormone classes and cell culture media components with a potential focus on added proteins like BSA.

Overall, our results demonstrate a strong direct effect of different phytohormone classes on *T. gondii* tachyzoite replication. To our knowledge, this is the first study to compare the effects of ABA, GIBB, and KIN in the same parasite culture model, and to investigate dose effects as well as the interaction with different cell culture media components typically used in *T. gondii*-infected host cell cultures.

Based on our study results, we suggest that ABA might be utilized to enhance tachyzoite replication in cell culture models for *in vitro* studies, while GIBB may potentially be pursued as potential *T. gondii* growth inhibitor based on the consistent decrease of tachyzoite numbers under the varied culture conditions. With GIBB being described as apoptosis enhancer (Sakr and Shalaby 2012), our results may indicate that GIBB is a promising agent for future testing as potential toxoplasmosis treatment. The use of both FBS and P/S additives is beneficial to *T. gondii* replication. Our results indicate that there is a strong impact of these cell culture media components, namely FBS and P/S, on the quality and quantity of phytohormone effects on *T. gondii* replication. While using these additives is not interfering strongly with the observations in ABA treated cultures, the effects of GIBB can be assessed more reliably without the additives. Further studies into underlying molecular mechanisms of general pathways triggered by different phytohormones seem crucial to elucidate the role of both phytohormones and their inhibitors in parasite growth. In addition, future investigations should focus on the impact of phytohormones on bradyzoites as these represent the natural infectious stage in tissue cysts.

## Acknowledgements

The authors would like to thank the personnel of the Institute of Parasitology at Leipzig University for their technical support and the animal husbandry. We would like to thank Lysanne Hiob and Martin Koethe for their gracious support in conducting the *in vitro* experiments.

## Declarations

### Ethical approval

No ethical approval was needed for this study, since all experiments were performed in vitro.

### Competing interests

The authors declare that they have no competing interests.

### Authors’ contributions

Tina Wagner (TW), Berit Bangoura (BB), and Arwid Daugschies (AD) designed the experimental setup. TW, Stefanie Wiedmer (SW), and BB conducted the experiments and compiled the data. TW, BB, and Ildiko Rita Dunay (IRD) analyzed the data and drafted the manuscript. All authors reviewed and approved of the manuscript.

### Funding

No external funding was provided for this study.

### Availability of data and materials

The original study data is with the authors.

## Notes

### Competing Interest Statement

The authors have declared no competing interest.

