## Supplementary material for "Phytohormones regulate asexual *Toxoplasma gondii* replication": Table

**Wagner et al. Phytohormones and their effect on asexual T. gondii stage development and replication - Tables**

**Table 1: Study design of pre-assay 2 (comparison of different *T. gondii*-infected MDBK culture conditions)**

| Experimental group (n=36) | *T.gondii* infection strain | cell culture medium and additives applied* |
| --- | --- | --- |
| ME49 | ME49 | IMDM |
| ME49F | ME49 | IMDM + 10 % FBS |
| ME49P | ME49 | IMDM + P/S |
| ME49FP | ME49 | IMDM + P/S + 10 % FBS |

* IMDM, Iscove’s modified Dulbecco’s medium; P/S, penicillin/streptomycin; FBS, fetal bovine serum

**Table 2: Study design of phytohormone efficacy experiments (all experimental groups of cell culture wells were infected with 5 x 10^5^ *T. gondii* tachyzoites immediately before the 2 h phytohormone treatment)**

| Experimental group | replicate number (n) | Phytohormone added^1^ | Phytohormone test concentration | cell culture medium used during treatment^2^ |
| --- | --- | --- | --- | --- |
| IUC | 36 | none | none | IMDM |
| A20ng | 12 | ABA | 20 ng/µl | IMDM |
| A2ng | 6 | ABA | 2 ng/µl | IMDM |
| A0.2ng | 6 | ABA | 0.2 ng/µl | IMDM |
| G250ng | 12 | GIBB | 250 ng/µl | IMDM |
| G50ng | 6 | GIBB | 50 ng/µl | IMDM |
| G10ng | 6 | GIBB | 10 ng/µl | IMDM |
| G2ng | 6 | GIBB | 2 ng/µl | IMDM |
| K50ng | 12 | KIN | 50 ng/µl | IMDM |
| K25ng | 6 | KIN | 25 ng/µl | IMDM |
| K1ng | 6 | KIN | 1 ng/µl | IMDM |
| IUC-F | 36 | none | none | IMDM + 10 % FBS |
| A20ng-F | 12 | ABA | 20 ng/µl | IMDM + 10 % FBS |
| A2 ng -F | 6 | ABA | 2 ng/µl | IMDM + 10 % FBS |
| A0.2 ng -F | 6 | ABA | 0.2 ng/µl | IMDM + 10 % FBS |
| G250ng-F | 12 | GIBB | 250 ng/µl | IMDM + 10 % FBS |
| G50ng-F | 6 | GIBB | 50 ng/µl | IMDM + 10 % FBS |
| G10ng-F | 6 | GIBB | 10 ng/µl | IMDM + 10 % FBS |
| G2ng-F | 6 | GIBB | 2 ng/µl | IMDM + 10 % FBS |
| K50ng-F | 12 | KIN | 50 ng/µl | IMDM + 10 % FBS |
| K25ng-F | 6 | KIN | 25 ng/µl | IMDM + 10 % FBS |
| K1ng-F | 6 | KIN | 1 ng/µl | IMDM + 10 % FBS |
| IUC-P | 36 | none | none | IMDM + P/S |
| A20ng-P | 12 | ABA | 20 ng/µl | IMDM + P/S |
| A2-Png | 12 | ABA | 2 ng/µl | IMDM + P/S |
| A0.2ng-P | 12 | ABA | 0.2 ng/µl | IMDM + P/S |
| G250ng-P | 12 | GIBB | 250 ng/µl | IMDM + P/S |
| G50ng-P | 12 | GIBB | 50 ng/µl | IMDM + P/S |
| G10ng-P | 12 | GIBB | 10 ng/µl | IMDM + P/S |
| G2ng-P | 12 | GIBB | 2 ng/µl | IMDM + P/S |
| K50ng-P | 12 | KIN | 50 ng/µl | IMDM + P/S |
| K25ng-P | 12 | KIN | 25 ng/µl | IMDM + P/S |
| K1ng-P | 12 | KIN | 1 ng/µl | IMDM + P/S |
| IUC-PF | 36 | none | none | IMDM + 10 % FBS + P/S |
| A20ng-PF | 12 | ABA | 20 ng/µl | IMDM + 10 % FBS + P/S |
| A2ng-PF | 12 | ABA | 2 ng/µl | IMDM + 10 % FBS + P/S |
| A0.2ng-PF | 12 | ABA | 0.2 ng/µl | IMDM + 10 % FBS + P/S |
| G250ng-PF | 12 | GIBB | 250 ng/µl | IMDM + 10 % FBS + P/S |
| G50ng-PF | 12 | GIBB | 50 ng/µl | IMDM + 10 % FBS + P/S |
| G10ng-PF | 12 | GIBB | 10 ng/µl | IMDM + 10 % FBS + P/S |
| G2ng-PF | 12 | GIBB | 2 ng/µl | IMDM + 10 % FBS + P/S |
| K50ng-PF | 12 | KIN | 50 ng/µl | IMDM + 10 % FBS + P/S |
| K25ng-PF | 12 | KIN | 25 ng/µl | IMDM + 10 % FBS + P/S |
| K1ng-PF | 12 | KIN | 1 ng/µl | IMDM + 10 % FBS + P/S |

^1^ ABA, abscisic acid, KIN, kinetin, GIBB, gibberellic acid

^2^ IMDM, Iscove’s modified Dulbecco’s medium; P/S, penicillin/streptomycin; FBS, fetal bovine serum; after the 2 h incubation period with the respective group-specific medium, all cultures were incubated for 48 h under identical culture conditions for parasite replication (IMDM + 10 % FBS + P/S)
